# Evaluation Of Biochemical, Hematological And Antioxidant Properties In Mice Exposed To A Triherbal (*Nigella sativa, Carica papaya and Boswellia sacra*) Formular

**DOI:** 10.1101/2020.08.02.233635

**Authors:** S. Kehinde, S. M. Adebayo, A. L. Adesiyan, E. A. Kade, K. Gurpreet

## Abstract

*Nigella sativa, Carica papaya and Boswellia sacra* are medicinal plants in the commonly used in folkloric medicine due to the presence of its immense therapeutic properties. Fifty (50) female albino mice weighing between 15-22g were divided into five groups of 10 mice each. Animal in group 1 served as control group and were administered distilled water while animal in group 2 were given 2ml of cisplatin (orally). Animal in group 3-5 were given orally; 100 mg/kg (low dose), 200 mg/kg (medium dose) and 400 mg/kg (high dose) of triherbal preparation. The feeding regimens lasted for 28 days. After 28 days, mammary gland and blood samples were collected for haematological and antioxidant analysis. The triherbal formula decreased the GSH and MDA levels of mice treated with 100 mg/kg and 400 mg/kg doses compare to control. The measurement of total protein content, SOD and CAT increased in treated animals compared to control. However, RBC (Red Blood Cell) counts significantly decreased in the low, medium and high dose groups (0.95±0.08, 6.57±0.08 and 3.55±0.55 x 10^6^ cells/mm^3^ respectively) compared to control (7.34±0.40) at P<0.05. Also, significant decreases (P<0.05) in the level of the total WBC (White Blood Cell) count, platelet count, PCV (Packed Cell Volume) and Hb (haemoglobin) concentration were observed. The decreases were dose dependent. The MCH (Mean Corpuscular Haemoglobin) and MCHC (Mean Corpuscular Haemoglobin Concentration) except MCV (Mean Corpuscular Volume) significantly decreased in treated group only. The triherbal formulation exhibited significant antioxidant activities showing increased levels of SOD, CAT and Protein content due to activation of the enzyme involve in detoxification of free radicals and decreased in the level of GSH and MDA due to accumulation of peroxides and H_2_O_2_. Also, decreased in haematological parameters due to the presence of phytochemicals such as phenol, resins, saponins, sterols, tannis and terpenes in the triherbal formula. Therefore, it has potential to induce haematotoxicity hence consumption of high concentrations should be discouraged.

## 1. Introduction

Over the years, plants have been used by humans as medicine to treat a vast number of diseases. The use of medicinal plants cuts across cultural lines as various traditional systems of medicine (Fabricant and Farnsworth, 2001). In Africa, the use of Egyptian traditional medicine dates from about 2900 B.C. In most African traditional societies, herbal remedies were often prepared as crude extract of medicinal plant organs such as leaves, roots, flowers and barks (Telefo *et al*., 2011; Fatima *et al.*, 2013).

Today, the popularity of traditional medicine has greatly increased across the world in both developed and developing nations. The World Health Organization estimates that about 80% of the populations in developing nations use traditional medicines, most of which are plant based remedies as complementary or alternative medicine (WHO, 2005).

Various factors can be attributed to the increase in the use of plant based remedies. They may include: economic considerations such as high cost of conventional medicines, perceived lower toxicity and fewer side effects of plant based medicines as these plants have been used before. To add on to the upsurge is the existence of diseases like cancer, to which no cure exists and the emergence of new diseases. The increased cases of drug resistance which are being encountered with the use of conventional medicines have favorably contributed to the use of plant based remedies (Bandaranayake, 2006; Abdullah, 2011; Pan *et al.*, 2014).

Plants have played an important role in drug discovery. For example vincristine and vinblastine which are used for the treatment of cancer are obtained from *Catharanthus roseus*. Quinine an antimalarial is obtained from *Cinchona ledgeriana* while digoxin is obtained from *Digitalis lanata* and is used as a cardiotonic (Fabricant and Farnsworth, 2001).

There are various ways through which plants can be used as sources of drugs. They include: using the whole plant or part of it as an herbal remedy, isolating bioactive compounds for direct use as therapeutic agents such as morphine. Plants can also provide raw materials for partial synthesis of drugs with higher activity or lower toxicity or they can be used as molecular models to produce new drugs (Fabricant and Farnsworth, 2001).

Despite the immense health benefits realized from use of plants as medicines, several challenges still exist such as insufficient scientific data to support use of some herbal remedies, lack of standardized formulation of herbal remedies and adulteration of herbal materials. According to the WHO, the assessment of the safety and efficacy of herbal remedies still remains a challenge (WHO, 2005; Ekor, 2014).

The use of medicinal plants is a practice among humans that has been passed down from one generation to another and plays a role in the development of human cultures and various traditional systems of medicine worldwide. According to the WHO, traditional medicine (TM) is defined as, “the sum total of knowledge, skills and practices based on the theories, beliefs and experiences indigenous to different cultures that are used to maintain health, as well as to prevent, diagnose, improve or treat physical and mental illnesses” (WHO, 2013). Based on fossil records, the use of medicinal plants dates back to the middle Paleolithic age 60000 years ago. These plants had a variety of uses such as food seasoning, weapons and medicines (Hassan, 2012).

Medicinal plants can be described as “any plant which, in one or more of its organs, contains substances that can be used for therapeutic purposes or as precursors for the synthesis of useful drugs”. The therapeutically useful phytochemicals obtained from plants include the alkaloids, flavonoids, tannins and phenolic compounds (Sofowora *et al.*, 2013; Choudhury *et al.*, 2015). In most plants, the quantity and the composition of bioactive compounds present are influenced by genotype, extraction procedure and environmental conditions (Dai and Mumper, 2010; Vinha *et al.*, 2011).

Plants are major part of most traditional medicine systems and a variety of conventional drugs have been obtained from plants following ethnobotanical leads from traditional remedies. Natural products and their derivatives represent over 50% of all drugs in clinical use worldwide according to Maridass and Britto (2008). In spite of these challenges, medicinal plants have a promising future to act as preventive medicine against various diseases and also as complementary medicine alongside conventional treatments so as to increase efficacy or reduce side effects of conventional therapies (Hassan, 2012). This study focused on establishing some medicinal plants used in treatment of cancer and also screen for their antioxidant activity and haematological parameters.

## 2. Materials and Methods

### 2.1. Plant materials and Sample preparations

Leaves of *Carica papaya* were sourced from Baale farmland, Asese, Obafemi Owode Government in Ogun State. The leaves were washed, air dried, and crushed to a powder with an electric micronizer. The black seeds and Frankincense were collected from the local markets. After that the seeds were grinded into fine powder form to prepare the crude alcoholic extracts. Two hundred gram of each of powdered plant material was kept in 1000ml of alcohol in conical flask. The mouth of the conical flasks were covered with aluminum foil and kept in a room temperature for 48 hours for complete elucidation of active materials to dissolve in ethanol. Then, the extracts were filtered by using muslin cloth followed by filter paper. The solvent form the extracts were removed with water bath at temperature of 40° C. Finally, the residues were collected and used for the experiment.

### 2.2 Animal Procurement and Conditioning

Fifty adult female mice were sourced from a local breeder at Otta in Ogun-State. The mice weighed between 14 g-25 g. They were kept in well ventilated cages cushioned with saw dust in the animal house of the Department Cell biology and Genetics, Faculty of science, University of Lagos. They were acclimatized for one week before actual experiment and kept under standard conditions of room temperature and 12:12 hours of light and dark cycle respectively. The mice were fed with standardized pellet and tap water ad libitum. The mice cages were regularly cleaned and saw dust changed every day.

### 2.3 Acute toxicity (LD_50_) study

A separate experiment was carried out to study the acute toxicity of the extracts on mice. Normal healthy female mice were randomly divided into 5 groups which fed with the vehicle-treated “control” groups (distill water) and three concentration of extract-treated “experimental” groups, totally making up to 5 groups of 10 animals per each group. Extract (50, 100, 200, 400 and 1000 mg/kg body weight) were orally administered to different test groups and control groups were separated. All the mice were allowed access to food and water. Behaviour changes and mortality were observed and recorded over a period of 72 hours. The LD50 was estimated from the graph of percentage (%) mortality (converted to probit) against log-dose of the extract, probit 5 being 50% (Aniagu et al., 2005).

### 2.4 Experimental Design and Grouping

The animals were divided in five groups of ten mice each. All mice were fed by normal diet and water ad-libitum. Mice in group A served as positive control, group B served as negative control, groups C, D, and E were administered by the alcoholic extracts once daily for a period of 28 days, with single dose of Cisplatin, 100, 200 and 400 mg/kg Body weight, respectively. All mice except from the negative control group were injected into the mammary fat with 0.1 mL of NMU. The mice were weighed three times a week and kept under normal temperature during the period of study.

**Table 2-1:**
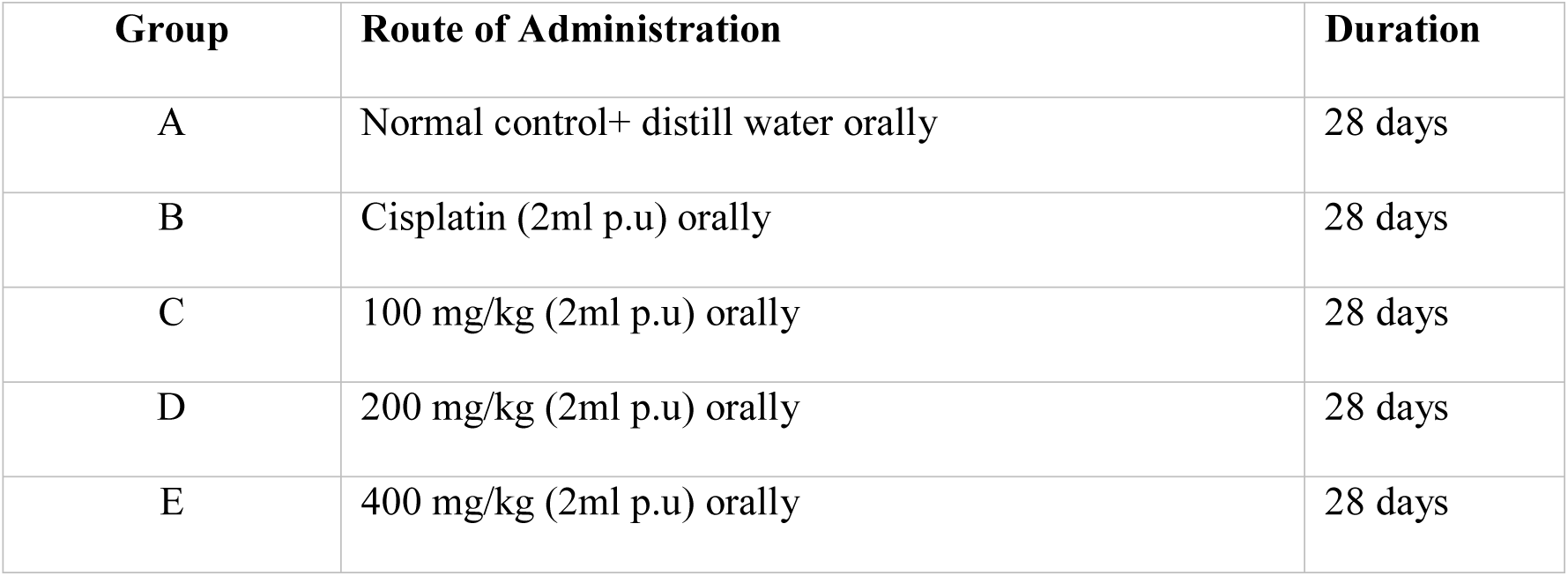
shows treatment and duration of groups.

#### Animals Sacrifice

The final body weight of the mice was obtained at the end of the treatment using a digital weighing balance. They were then sacrificed by decapitation twenty four hours after the last treatment. Blood samples were collected and taken in EDTA containing tubes from animals of different groups for haematological measurements. Moreover, mammary tissues were fixed for antioxidant investigation.

#### Ethical Approval

The study was conducted in accordance with the declaration of Helsinki and was approved by the local institutional review committee and the Health Research Ethics Committee (HREC) of Lagos University Teaching Hospital (LUTH) with HREC assigned number ADM/DCST/HREC/APP/854

### 2.5 Haematological Measurements

Complete blood count (CBC) includes hemoglobin content, red blood cells (RBC), white blood cells (WBC), was done by using Automated Hematology Analyzer, ready–made kits and platelets (PLT) counts.

#### 2.5.1 Determination of packed cell volume (PCV)

The blood in the EDTA bottle was used for the PVC. The blood was collected into a capillary tube containing anticoagulant. Plug one end of the tube with soft wax to a depth of about 2mm by heating it carefully over a flame. Place the capillary tube in the numbered slots in heaematocrit centrifuge. After centrifuge at high speed (13000 rpm) for 5 minutes. The percentage of PVC is determined using haematocrits was calculated based on the following formula

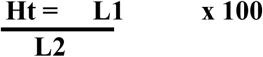

Where,

Li = is the height of RBC column

L2 = is the total length of the column (RBC + Plasma + buffy coat) in millimeter and expressed in per cent

#### 2.5.2 Determination of total white blood cell counts

The counting of total white blood cells was done by using a diluting fluid (Turk’s fluid) in a ratio of 1:20 which haemolyses the RBCs leaving the WBCs to be counted. The leukocytes are the counted in a counting chamber under the microscope, and the number of cells in a litre of blood is calculated.

#### 2.5.3 Determination of heamoglobin (Hb)

Sahli’s haemoglobinometer was employed for estimation of haemoglobin (Hb) content of the blood. Shahi’s pipette was filled with mice blood exactly up to 20 mm^3^ mark. The excess of blood was removed by blotting the tip with soft absorbent tissue. The blood was expelled into a calibrated (transmission) test tube containing 1 ml of 0.1 N HCl acid solutions and the pipette was rinsed several times in the acid solution. The sample was allowed to stand for 3 minutes. This method involves conversion of hemoglobin to acid haematin. The amount of haemoglobin in the blood sample was directly read in gram percent from the graduated haemoglohinometer tube.

**Figure 1:**
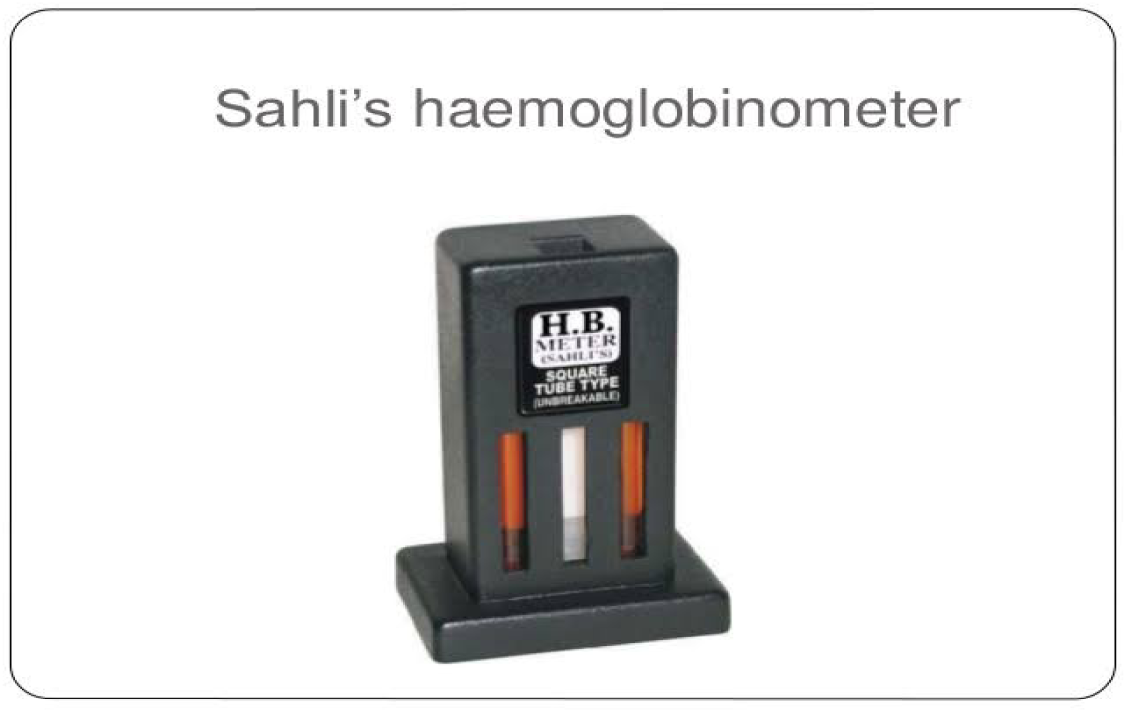
Sahli’s haemoglobinometer

#### 2.5.4. Other blood indices

Haematological indices such as Mean Corpuscular Volume (MCV), Mean Corpuscular Haemoglobin Concentration (MCHC) and Mean Corpuscular Haemoglobin (MCH) were calculated from the values of Hh content (%) and Ht (%) using the following formula

a. 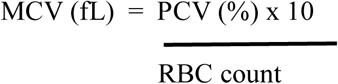
b. 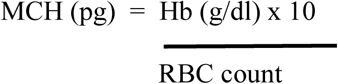
c. 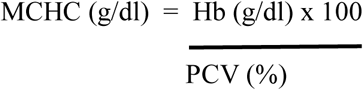

#### 2. 6. 5. Differential blood counts (DC)

The differential counting was done as described in clinical haematology. The blood smears were made, air-dried, fixed in 100% methanol and stained with May and Grunwald stain and counted under oil immersion objective. Smears were examined for macrophages and abnormal RBC morphology (size, shape, colour, maturity, inclusions) and to determine the differential count of white blood cells (WBC). Total of 1000 blood cells of all types was counted from each smear and then percentage of each cell type was calculated.

##### 2.5.5.1 May-Grünwald staining

- Since the May-Grünwald staining solution is made up in MeOH prior fixation is not necessary.
- Place slide on a flat surface and pipet 500 μl May-Grünwald Stain on the slide, leave for 3 min.
- Dilute Stain by adding 500 μl 10mM NaPi 7.0, leave for 7 min.
- Lift slide to drain the staining solution and place in a tray with H_2_O for 1 min.
- Dry slide vertically for 5 min.
- Mount coverslips using an aqueous-based mounting medium.

### 2.6. Biochemical Analyses

#### 2.6.1. Sample preparation (tissue homogenate)

Breast tissues were collected from above groups and processed. Breast tissue was perfused with saline to remove any red blood cells and clots. Tissue was homogenized with the saline (0.9%) (1 g breast in 10 ml saline) with ice-cold PBS pH 8.0 using a homogenizer (Yamato LSC LH-21, Japan) and centrifuged at 12,000 rpm for 30 min at 4°C. Supernatant was collected and used for following biochemical estimations.

#### 2.6.2. Protein estimation

Total protein contents were estimated by the modified method of Lowry *et al*. (1951). 0.5 ml of homogenized tissue is mixed with 1.5 ml of 0.2 M Tris buffer (pH-8.2) and 0.1 ml of 0.01 M DTNB and this mixture is brought to 10.0 ml with 7.9 ml of absolute methanol. The above reaction mixture is centrifuged at approximately 300 g at room temperature for 15 minutes. The absorbance of supernatant is read in a spectrophotometer against reagent blank (without sample) at 412 nm. Tissue protein is than calculated with reference to the standard graph and the results were expressed as milligram protein per gram of tissue weight.

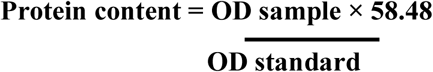

Where;

**OD** = Optical density at 412 nm

#### 2.6.3. Estimation of glutathione

Glutathione (GSH) contents were measured as total non-protein sulfhydryl (NPSH) group using the method of Moron *et al*. (1979) with modifications. For the measurement of GSH content, 1.6 ml sodium phosphate buffer, 0.1 ml of 1 mM ethylenediamine tetra acetic acid disodium salt (EDTA, Amresco), 0.1 ml nicotinamide adenine dinucleotide phosphate reduced (NADPH) and 0.1 ml oxidized glutathione as well as PMS (0.1ml) in total volume of 2ml. The enzyme activity is measured at 340 nm and calculated as nanomole NADPH oxidized/min/mg of protein using extinction coefficient of 1.36 × 10^3^ M/cm. The change in absorbance/min was determined and this value was converted to micromole GSH in comparison to a known standard.

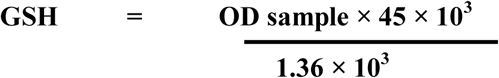

Where;

**OD** = Optical density at 340 nm

**1.36 × 10**^**3**^ = Extinction coefficient

#### 2.6.4. Estimation of Superoxide Dismutase Activity (SOD)

Superoxide dismutase (SOD) activity was assayed by the nitroblue tetrazolium (NBT) method as described by Beauchamp *et al*. (1971). In this method, the reaction mixture consists of 0.5ml supernatant, 1ml of 50mM Sodium carbonate, 0.4ml of 25μM NBT, 0.2ml of 0.1mM EDTA. The reaction is then initiated by the addition of 0.4ml of 1mM hydroxylamine hydrochloride. The change in absorbance is recorded at 560 nm using a UV spectrophotometer. The control is simultaneously run without homogenate. Units of SOD activity are expressed as the amount of enzyme required to inhibit the reduction of NBT by 50 %. Specific activity of total SOD is expressed as units per milligram protein.

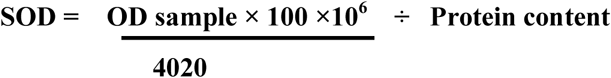

#### 2.6.6. Estimation of catalase in breast

Catalase (CAT) activity was determined by catalytic reduction of hydrogen peroxide using a standard method described by Aebi (1984). The mixture consists of 1.95 ml of phosphate buffer (0.05 M, pH-7), 1 ml of H2O2 (0.019 M) and 0.05 ml sample (10 % w/v) in a final volume of 3 ml. control cuvette contains all the components except substrate. Change in absorbance is then recorded at 240 nm and the results were expressed as micromole of product formed per minute per milligram protein of the tissue.

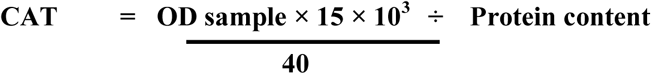

#### 2.6.7. Estimation of Malondialdehyde Level in breast

MDA levels, an index of lipid peroxidation were measured by double heating method of Okhawa *et al*, (1979). The method is based on spectrophotometric measurement of the purple colour generated by the reaction of TBA with MDA. For this purpose, 2.5 mL of trichloroacetic acid solution (10%w/v) was added to 0.5mL homogenized tissue in each centrifuge tube; the tubes were then placed in a boiling water bath for 15mins. After cooling to room temperature, the tubes were centrifuged at 1000xg for 10mins and 2mL of each sample supernatant was transferred to attest tube containing 1 mL of TBA solution (0.67% w/v). Each tube was then placed in a boiling water bath for 15min. After cooling at room temperature, the absorbance was measured at 532 nm by using spectrophotometer. The concentration of MDA was calculated based on absorbance coefficient of the MDA complex (e= 1.56×10^5^ cmM^−1^).

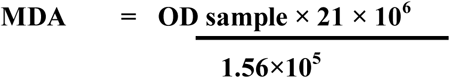

Where;

**OD** = Optical density at 532 nm

**1.56×10**^**5**^= Extinction coefficient

### 2.7 Statistical analysis

Experimental results are expressed as mean ± standard error of the mean (mean±S.E.M). The data were analysed by ANOVA (p<0.05) and means separated by Duncan’s multiple range tests (by SPSS version 21 software). Tabulation and graphics of data were done using Microsoft Excel XP.

## 3. Result

### 3.1. Morphological results

Table 1 demonstrates the changes in the body weight of mice after induction of NMU and during the periods of treatment with extracts. There was a significant difference at (p<0.05) between the treatment groups and normal control group, which signifies the extracts increases the weight of the animals. Figure 3-1 illustrates that the weight between all Alcoholic extracts-treated groups and controls were significantly different (P > 0.05).

**Figure 3-1:**
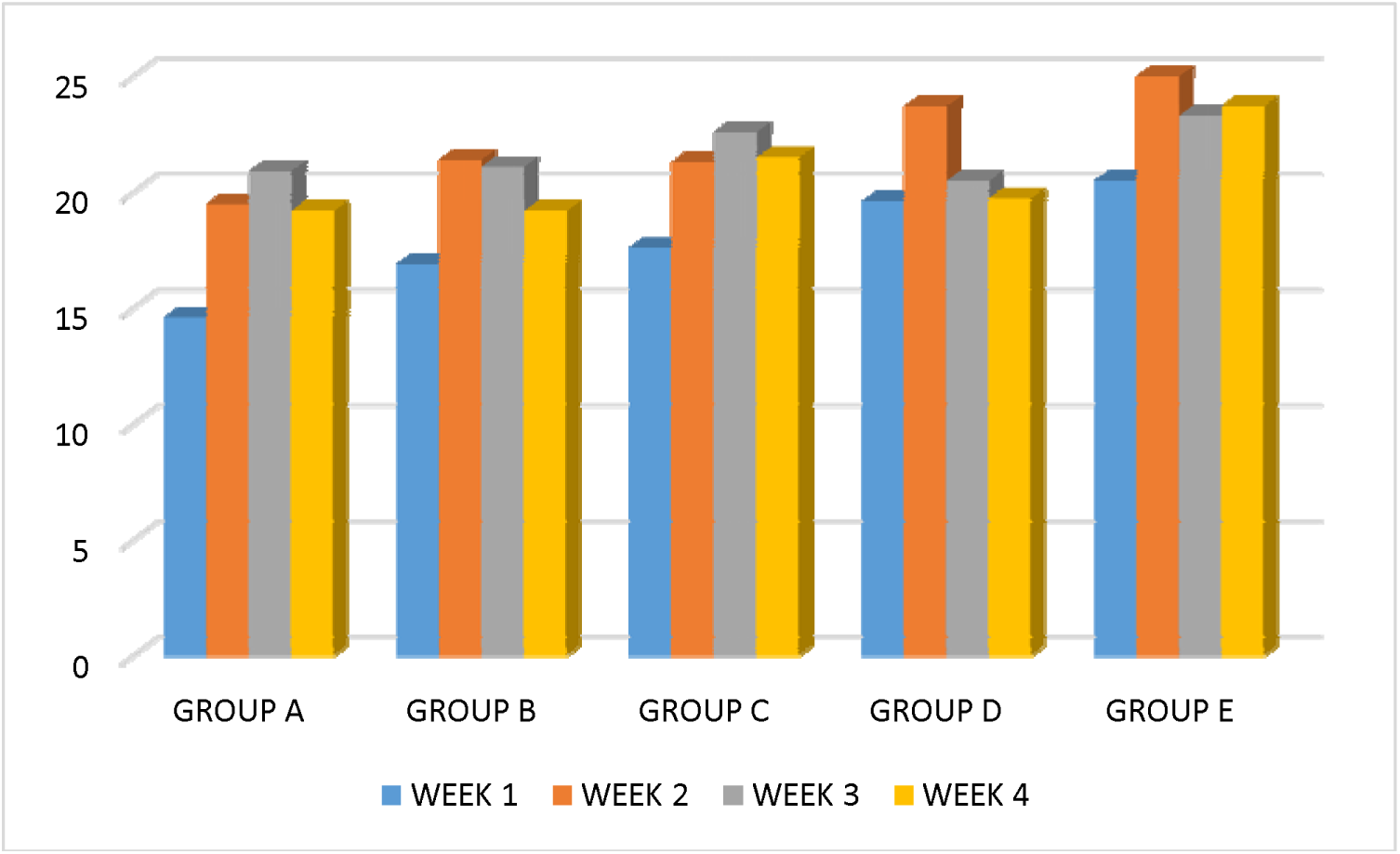
Change in the body weight mice treated with dose of alcoholic extracts

In Table 3-1. Results expressed as mean ± S.E.M of the mean body weight of female mice during the experiment in grams

**Table 3-1.** Mean initial and final body weight of adult female mice.

| Groups | Initial weight (g) | Final weight (g) |
| --- | --- | --- |
| **A (Normal control)** | 14.6 ± 0.37 | 19.2 ± 1.01 |
| **B (Cisplatin)** | 16.9 ± 0.40 | 19.6 ± 0.55 |
| **C (100 mg/kg)** | 17.6 ± 0.16 | 21.5 ± 1.04 |
| **D (200 mg/kg)** | 19.6 ± 0.16 | 19.8 ± 0.85 |
| **E (400mg/kg)** | $20.5 \pm 0.26$ | $23.7 \pm 0.88$ |

#### 3.1.1. Organ to body weight ratio

The organ to body weight ratios of Alcoholic extracts -treated groups and controls are illustrated in Table 3-2. The treatment groups (100, 200 and 400 mg/kg of extracts) and the positive control showed significant increase of lung, heart and liver to body weight ratio (P < 0.05) compared to the negative control. The liver to body weight ratio of the 400 mg/kg Alcoholic extracts-treated group decreased significantly (P < 0.05) compared to the positive control.

In Table 2, results expressed as Mean ± Standard Error Mean (S.E.M) of the mean organs to body weight of the mice during the experiment in grams. Values were significantly different (p< 0.05).

**Table 2.** Mean organs to body weight ratio of adult mice weight

|  | **LIVER** | **RT.KIDNEY** | **LT. KIDNEY** | **HEART** | **LUNG** |
| --- | --- | --- | --- | --- | --- |
| **A(control)** | $0.99 \pm 0.03$ | $0.1 \pm 0$ | $0.1 \pm 0$ | $0.1 \pm 0$ | $0.16 \pm 0.02$ |
| **B**<br>**(Cisplatin)** | 1.31±0.02 | 0.1±0.0 | 0.1±0.0 | 0.1±0.0 | 0.15±0.02 |
| **C (100**<br>**mg/kg)** | 1.1±0.0 | 0.1±0.0 | 0.1±0.0 | 0.1±0.0 | 0.3±0.0 |
| **D (200**<br>**mg/kg)** | 1.5±0.05 | 0.1±0.0 | 0.1±0.0 | 0.1±0.0 | 0.13±0.33 |
| **E**<br>**(400mg/kg)** | 0.27±0.21 | 0.1±0.0 | 0.1±0.0 | 0.2±0.0 | 0.17±0.06 |

**Figure 3-2.**
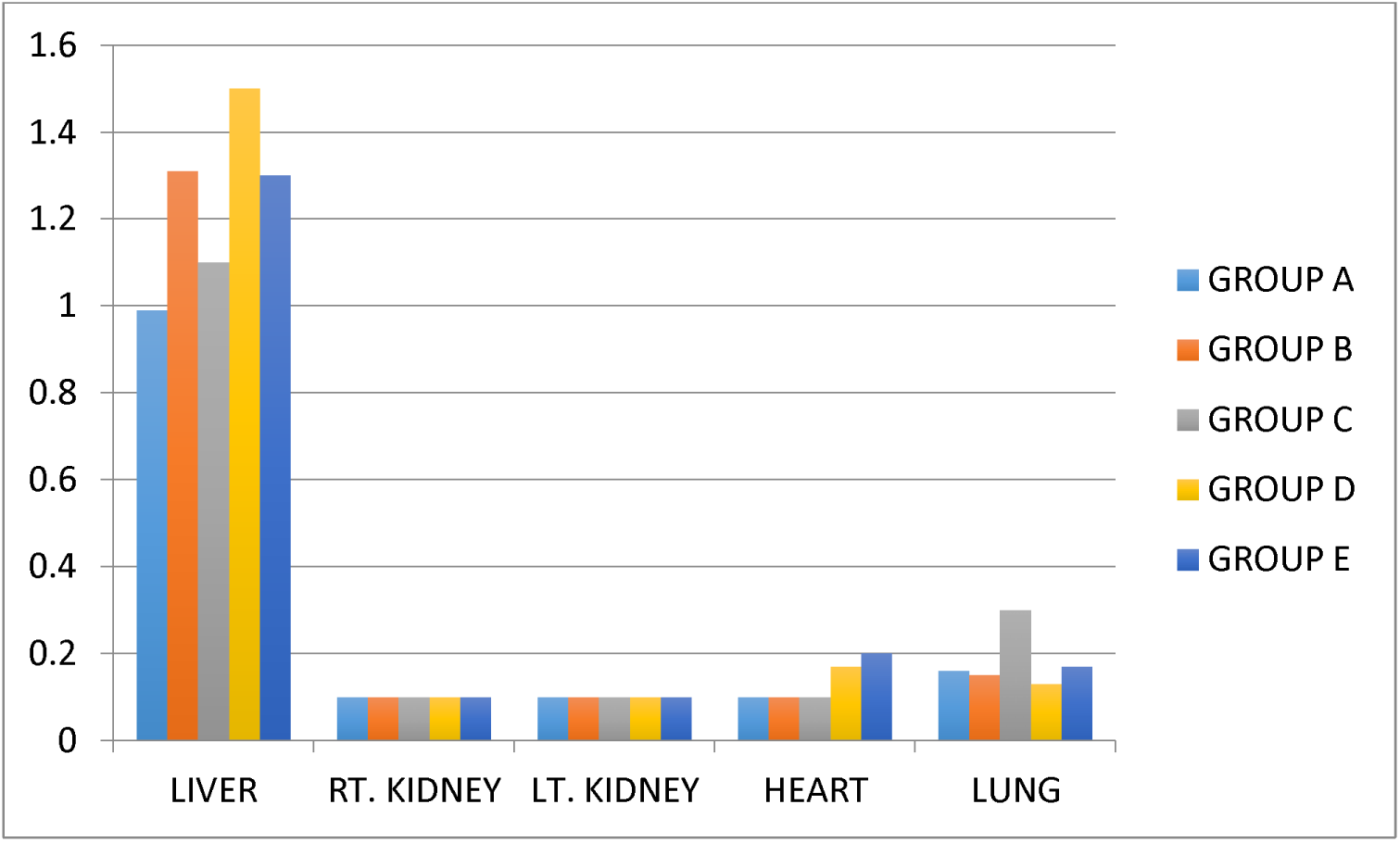
Organ to body weight ratio mice treated with extract. Mean were significantly different (P < 0.05).

### 4.2. Antioxidant biomarkers result

Table 3 shows results obtained from the evaluation of selected antioxidants biomarkers of breast tissues of experimental mice. There is no significant difference (P>0.05) in the value obtained from catalase activity, superoxide dismutase and total protein when compare to the control groups, however, glutathione and malondialdehyde showed significant difference p<0.05 at plant concentration of 100mg/kg, 200mg/kg and 400mg/kg respectively. There is also a significant difference in the superoxide dismutase values of the cisplatin group and control group. The levels in the antioxidant parameters indicating biomarkers of mammary gland are illustrated in Figure 3.

**Table 3:** Comparison of selected antioxidants biomarker of mammary gland of experimental mice.

| S/N | Antioxidant Biomarkers | Control | Cisplatin | 100mg/kg | 200mg/kg | 400mg/kg |
| --- | --- | --- | --- | --- | --- | --- |
| 1 | Catalase | 1.01±0.00 | 1.44±0.01 | *1.02±0.00 | *0.98±0.01 | *0.96±0.01 |
| 2 | Superoxide Dismutase | 12.5±0.36 | 9.10±0.51 | 14.0±0.38* | 9.56±0.38 | *14.0±0.26 |
| 3 | Glutathione | 4.00±0.19 | 4.57±0.02 | *3.8±0.02 | *3.67±0.02 | *3.56±0.03 |
| 4 | Malondaildehyde | 24.14±0.11 | 25.31±0.08 | 5.42±0.12 | 13.06±0.16 | 6.64±0.12 |
| 5 | Total protein | 43.55±0.15 | 45.56±0.20 | *44.4±0.20 | *45.95±0.13 | *44.4±0.20 |

Values are means of 3 replicates ± Standard Error of the Mean (S.E.M) and Values carrying superscript (\*) Non-significant between control groups and animal treated with dose of (100mg/Kg, 200mg/Kg and 400mg/Kg) of alcoholic extract

Values are means of 3 replicates ± Standard Error of the Mean (S.E.M) and Values carrying superscript (*) Non-significant between control groups and animal treated with dose of (100mg/Kg, 200mg/Kg and 400mg/Kg) of alcoholic extract

**Figure 3-3:**
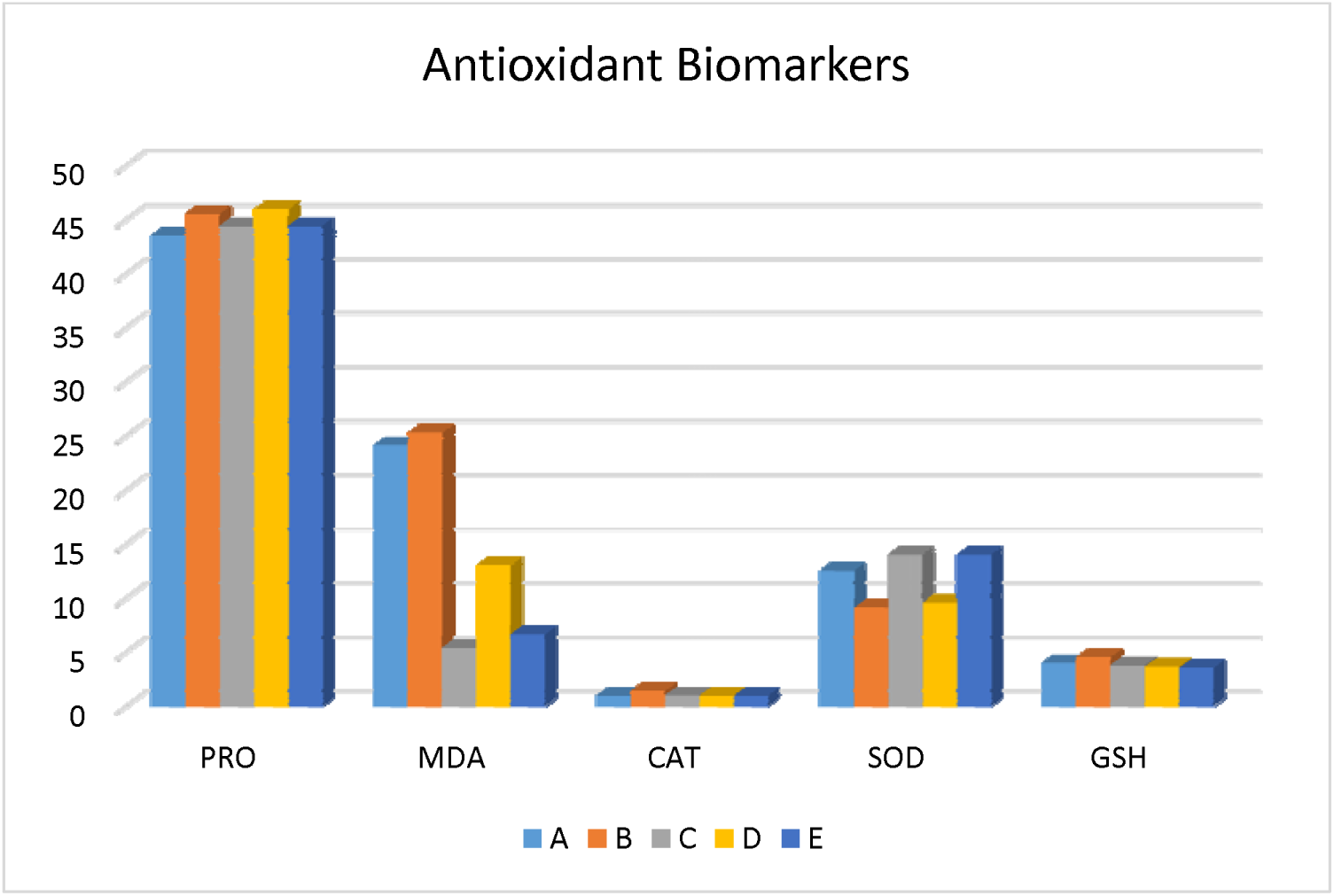
Antioxidant profile in control and experimental female mice

### 3.3 Effect of plants extract on the Haematological parameters

Studying the haematological parameters revealed that there is a significant (p≤0.05) decrease in white blood counts (WBCs), red blood counts (RBCs), Platelets count (PLC) and counts in addition to haemoglobin count after administration of 100 mg/Kg, 200 mg/Kg and 400 mg/Kg body weight, respectively, while the dose of 100 mg/Kg body weight induced changes when compare with normal control group. Moreover, none of these doses cause any change in the platelet count as shown in Table 4. Comparing the values of the treated groups were significantly effective when compared with 100mg/Kg treated one (p<0.05) for RBCs. 200 mg/Kg treated group showed appreciated Hb content when compare with 100mg/Kg and 400 mg/Kg treated ones (Figure 4).

The mean PCV in control group was 35.00±0.00% while those of 100mg/kg, 200mg/kg and 400mg/kg dose groups were 34.5±0.50%, 33.5±2.0% and 32.00±1.00% respectively. The mean PCV of the 100mg/kg and 400mg/kg dose group were significantly different from that of control group (P<0.05) while the medium dose group was not significantly different. Also, the mean Hb (Haemoglobin) concentration in 100 mg/kg (2.0±0.30g/dl) and 400 mg/kg dose (5.65±0.15g/dl) groups were statistically significant compared with control group (12.25±0.15g/dl) while that of 200 mg/kg dose group (12.05±0.10g/dl) did not differ from control values. The mean platelet count of 100 mg/kg (223.00±7.00 x 10^3^ cells/mm^3^), 200 mg/kg (605±11.00 x 10^3^ cells/mm^3^) and 400 mg/kg dose (399±2.50 x 10^3^ cells/mm^3^) groups were significantly different compared with that of control group (920.00±247 x 10^3^ cells/mm^3^). The mean values of MCV for the control, 100 mg/kg, 200 mg/kg and 400 mg/kg dose groups were 48.00±3.00, 49.50±1.50, 50.50±1.50 and 47.00±2.001fL respectively. These values were not significantly different from each other. The mean values of MCH were also not significantly different among the groups when compared with the control group (18.50±0.50pg). Also, the 100 mg/kg (38.90±1.00g/dl), 200 mg/kg (33.00±1.00g/dl) and 400 mg/kg (39.00±1.50g/dl) dose groups of MCHC were significant different compared with the control group (39.50±1.00g/dl, P>0.05).

Result expressed as Mean ± SEM. ANOVA (p value) represents the difference between all groups. (*) Non significant between control groups and animal treated with dose of (100mg/Kg, 200mg/Kg and 400mg/Kg) of alcoholic extract.

**Table 3-4:** Effect of oral administration of daily doses of Alcoholic extract on haematological parameters of normal female mice.

| PARAMETERS | Control | Cisplatin | 100mg/kg | 200mg/kg | 400mg/kg |
| --- | --- | --- | --- | --- | --- |
| Red Blood cell | $7.34\pm0.40$ | $6.57\pm0.37$ | $0.95\pm0.08$ | $*6.57\pm0.08$ | $3.55\pm0.55$ |
| Haemoglobin | $12.25\pm0.15$ | $12.05\pm0.05$ | $2.0\pm0.30$ | $*12.05\pm0.10$ | $5.65\pm0.15$ |
| PVC | $35.50\pm0.0$ | $34.5\pm0.50$ | $*33.5\pm0.50$ | $*34.5\pm2.0$ | $*32.00\pm0.50$ |
| MCV | $48.00\pm3.0$ | $49.5\pm1.50$ | $*50.5\pm1.50$ | $*49.5\pm2.0$ | $*47\pm2.00$ |
| MCH | $18.50\pm0.50$ | $17.00\pm0.0$ | $*17.5\pm0.50$ | $*17.00\pm1.0$ | $*17.5\pm0.50$ |
| **MCHC** | 38.00±1.00 | 33.00±1.0 | *38.00±1.0 | 33.00±1.0 | *39.5±1.50 |
| **RDW** | 12.9±0.70 | 11.85±0.75 | *11.75±0.25 | *11.85±0.10 | *11.3±0.20 |
| **White Cell count** | 4.35±1.15 | 3.4±0.20 | *3.85±0.15 | *3.4±0.10 | *4.15±0.15 |
| **Neutrophils** | 0.76±0.29 | 0.5±0.04 | *0.11±0.01 | *0.5±0.02 | *0.09±0.01 |
| **Lymphocytes** | 4.02±0.58 | 2.72±0.28 | *3.83±0.06 | *2.72±0.11 | *3.02±0.01 |
| **Monocytes** | 0.05±0.03 | 0.025±0.01 | *0.015±0.05 | *0.025±0.00 | *0.015±0.01 |
| **Eosinophile** | 0.18±0.06 | 0.15±0.03 | *0.02±0.05 | *0.15±0.00 | *0.02±0.01 |
| **Basophils** | 0.37±0.23 | 0.29±0.16 | *0.11±0.07 | *0.29±0.01 | *0.18±0.01 |
| **Platelets count** | 920±247 | 905±262 | 223±7.0 | 605±11 | 399±2.50 |

Result expressed as Mean ± SEM. ANOVA (p value) represents the difference between all groups. (\*) Non significant between control groups and animal treated with dose of (100mg/Kg, 200mg/Kg and 400mg/Kg) of alcoholic extract.

**Figure 3-4:**
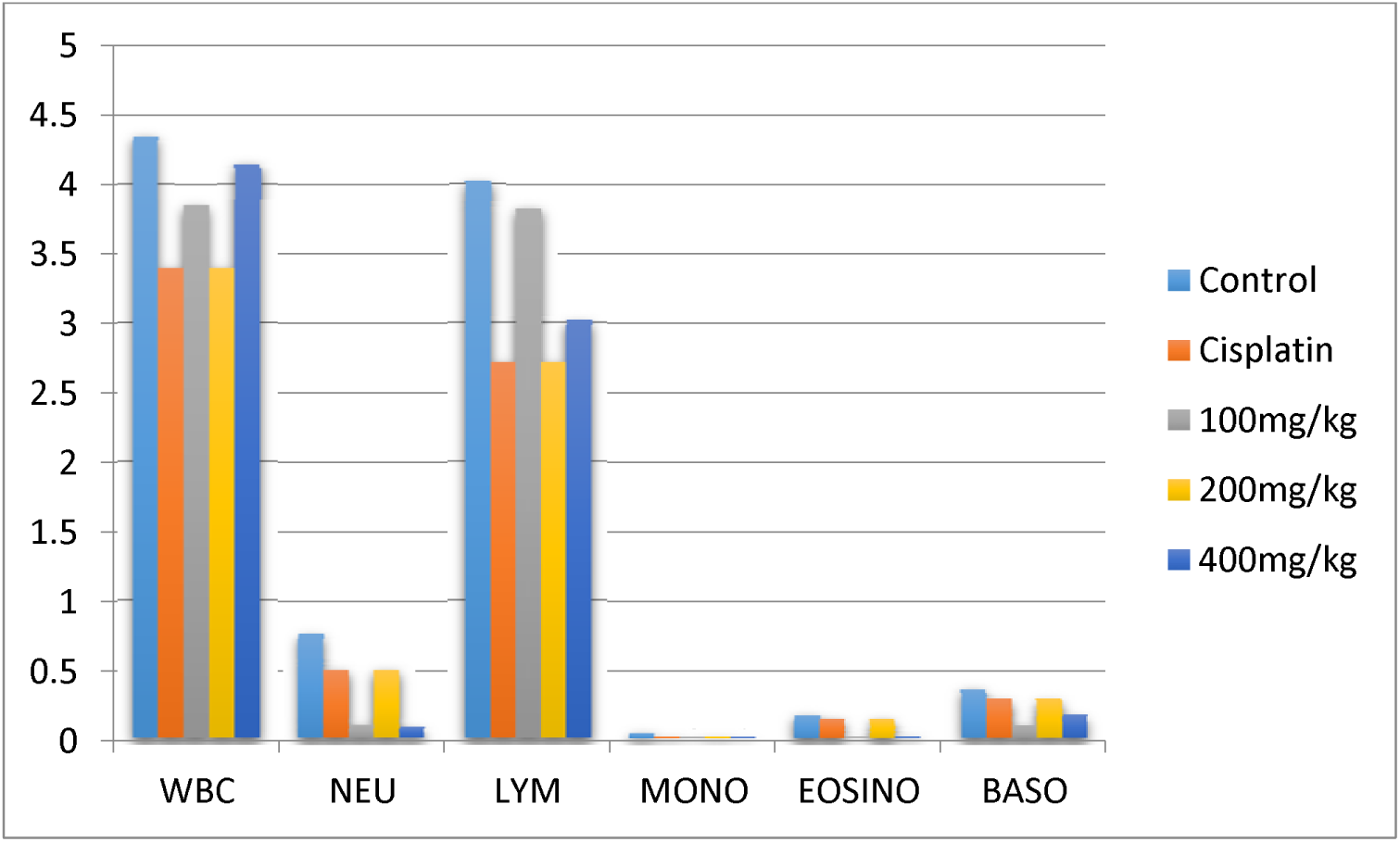
Effect of plants extract on the Haematological parameters

**Figure 3-5:**
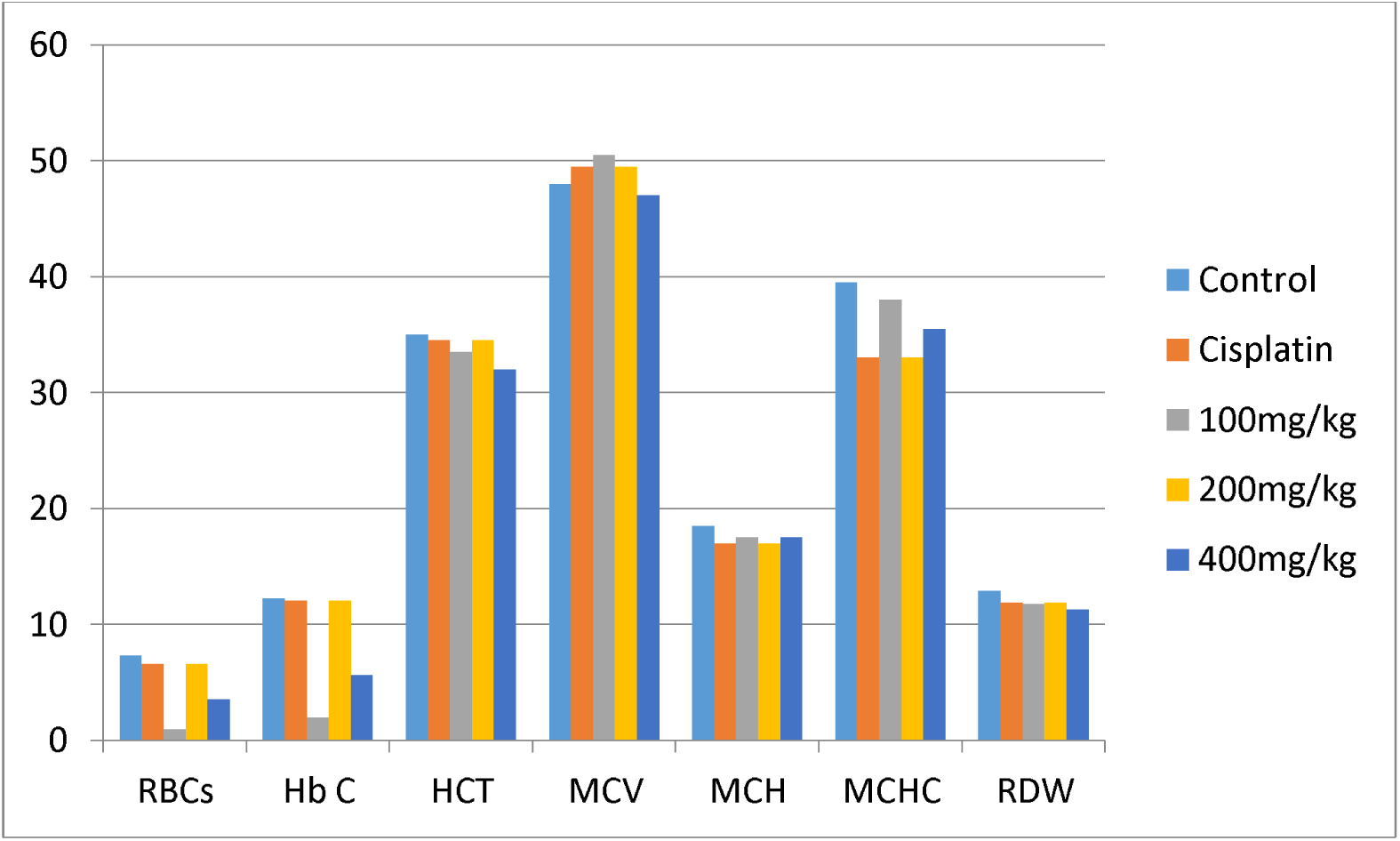
Effect of plants extract on the RBCs, WBCs and Hemoglobin content

**Figure 3-6:**
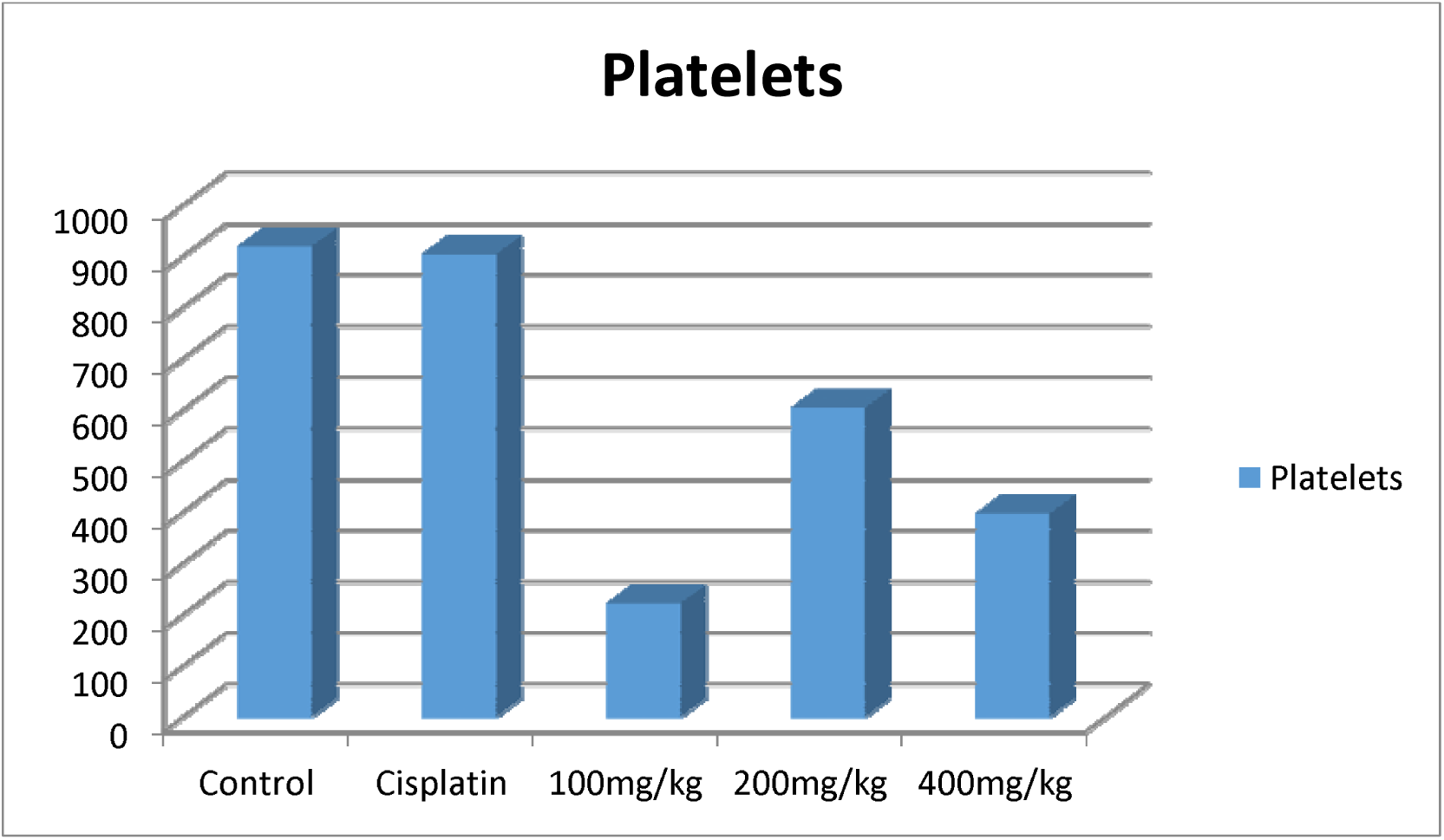
Effect of Alcoholic extracts oral treatment on alteration of Platelets count

## 4. Discussion

During this study, the routine weight gained over the period of exposure may be due to the presence of some phytochemicals in the extract. Tannins have been previously implicated in increasing body mass (Marcus *et al.*, 2003). Decreased GSH content observed in this study indicated impairment in cell’s defense against ROS and has been known to cause cellular injury (Omoreigie and Osagie, 2011) and generally reflects the inability of a tissue to scavenge excess superoxide anions leading to oxidative stress (Omoreigie and Osagie, 2007; Shafaquat *et al.*, 2017). MDA levels decreased in a dose dependent manner in the breast tissue. A significant MDA decrease was observed due to lipid peroxidation which is a direct indicator that cell membrane damage has occurred in the tissue (Jonas *et al.*, 2000; AshokKumar, 2004). Increased CAT observed in this study may indicate enhanced triherbal toleration by that particular tissue (Bahrami *et al.*, 2015). Rise in SOD activities observed may indicate presence of active enzyme involvement in neutralizing the effect of free radicals (Deger *et al.*, 2008). Elevated levels of protein content were noticed in all treated mice which may imply that the cell is capable of mitigating effect of free radical and peroxide processes which could ultimately results in modulating the host antioxidant status (Siwela *et al.*, 2013).

The haematological studies of triherbal preparation showed severe anaemia, which may imply inhibition of globin synthesis, depression of erythropoiesis, or a decreased level of folic acid (Antai *et al.*, 2009; Atasaya *et al.*, 2009; Yadav *et al.*, 2010). Extract administration might have caused destruction of erythrocytes directly or the decreased RBC count may be due to the effect of extract on erythropoietic tissue (Antai *et al.*, 2009). The manifestation of hypochromic anaemia is due to reduction in the number of red blood cells or haemoglobin or impaired production of erythrocytes (Antai *et al.*, 2009; Chia *et al.*, 2009). Combine extract might be responsible for the decreased RBCs and haemoglobin levels due to increased level of pro-inflammatory cytokines that induced iron retention by reticulo-endothelial system, gastrointestinal tract and liver, thereby exerting inhibitory effect on erythroid precursors (Harnafi and Amrani, 2007). The significant decrease in WBC observed in this study may be alluded to suppression of the haematopoetic system, which consequently reduces the production of WBCs (Ekiz *et al.*, 2005), and bio concentration of the toxicant in the kidney and liver (Amusa *et al.*, 2003). Also, decreased level of white blood cell counts were observed mainly in mice exposed to extract due to the fact that triherbal formula may induce leucopenia and thrombocytopenia in cases of severe liver dysfunction (George, 2000) and as a result of decreased defence mechanism against probable attack of toxic molecules during extract toxicosis (Kori-Siakpere, 2011). Decreased in haematocrit observed in this study can be attributed to the reduction in RBC count caused by either destruction or reduction in size (Schneider *et al*, 2003).

Variation in MCV, MCH, and MCHC values observed in this study may imply that the macrocytic anaemia which can lead to very slow production of erythroblasts in bone marrow (Ghaffar *et al.*, 2014) which make them grow over in size with shape and have fragile membranes called megaloblast which is characteristic of pernicious anaemia which can lead to megaloblast anaemia (Hussain *et al.*, 2014). The reduction in Hb, RBC, WBC, MCV, MCH, and MCHC indicated that there is slow development of blood in the haemopoitic cells (Sharaf *et al*., 2010) due to the presence of saponin in the tri-herbal preparation which has been reported to as reported to suppress haematopoiesis of all blood cells (Akinnuga *et al.*, 2011).

In conclusion, the tri-herbal formulations at doses evaluated has potential to induce haematotoxicity and indiscriminate consumption of high concentrations should be discouraged. Although these medicinal plants may possess profound therapeutic advantages at very low doses. Further research should be carried out in lower doses to ascertain the safety.

